# Explainability methods for differential gene analysis of single cell RNA-seq clustering models

**DOI:** 10.1101/2021.11.15.468416

**Authors:** Madalina Ciortan, Matthieu Defrance

## Abstract

Single-cell RNA sequencing (scRNA-seq) produces transcriptomic profiling for individual cells. Due to the lack of cell-class annotations, scRNA-seq is routinely analyzed with unsupervised clustering methods. Because these methods are typically limited to producing clustering predictions (that is, assignment of cells to clusters of similar cells), numerous model agnostic differential expression (DE) libraries have been proposed to identify the genes expressed differently in the detected clusters, as needed in the downstream analysis. In parallel, the advancements in neural networks (NN) brought several model-specific explainability methods to identify salient features based on gradients, eliminating the need for external models.

We propose a comprehensive study to compare the performance of dedicated DE methods, with that of explainability methods typically used in machine learning, both model agnostic (such as SHAP, permutation importance) and model-specific (such as NN gradient-based methods). The DE analysis is performed on the results of 3 state-of-the-art clustering methods based on NNs. Our results on 36 simulated datasets indicate that all analyzed DE methods have limited agreement between them and with ground-truth genes. The gradients method outperforms the traditional DE methods, which encourages the development of NN-based clustering methods to provide an out-of-the-box DE capability. Employing DE methods on the input data preprocessed by clustering method outperforms the traditional approach of using the original count data, albeit still performing worse than gradient-based methods.

## 1 Scientific Background

The advancements in single-cell sequencing (scRNA-seq) motivated the proposal of numerous methods for a dedicated computational analysis. These efforts are motivated by the specific scRNA-seq technical challenges, residing in the high data sparsity, accentuated by dropout, which in this context represents a technical noise corrupting the in the expression matrix with false zero values. scRNA-seq data consists of a count matrix *D* = {*xij*} ∈R^*d*\**n*^ with *d* features (i.e. genes) and *n* samples (i.e. cells). As cell annotations are typically lacking, clustering methods are routinely employed to find groups of cells representing distinctive cellular subtypes. After the cellular clusters are detected, the molecular analysis of the phenotypic variation studies the quantitative differences between clusters or conditions. To this end, multiple differential expression analysis (DE) methods have been proposed to identify the genes expressed differently across detected clusters. The first DE analysis methods consisted of statistical tests (such as t-test, Wilcoxon). Several methods proposed for the DE analysis of bulk RNA-seq are also employed on scRNA-seq. For example, edgeR estimates a common dispersion for all isoforms using conditional maximum likelihood and proposes two hypothesis tests for the DE: a quasi-likelihood F-test and likelihood ratio test. Additionally, multiple dedicated scRNA-seq methods were also proposed. DEsingle estimates the proportion of real and dropout zeros using a zero-inflated negative binomial regression model and then classifies the DE genes into three categories.

However, as reported by several broad comparative studies [1] [2], there is no consensus on the best performing method and the agreement between DE methods is generally very low. In [3], 11 DE methods were shown to have 92 genes in common from top 1000 and only 41 overlapped with the gold standard. A set of 9 tools for DE analysis was analyzed in [1], confirming the same poor agreement and profiling the studied methods when the count data has varying levels of expression. The DE methods presented so far are model agnostic: they build a standalone model on the input data which can be applied to the result of any prior clustering to identify the underlying DE genes.

In parallel, as explainability became a common requirement in machine learning (ML), model-agnostic methods became adopted in various fields to make the decisions of predictive models more transparent and understandable. Despite the fact that explainability and interpretability are often used interchangeably, works like [5] clarify the distinction. Explainability provides post hoc explanations for existing black box models while interpretability studies models that are inherently interpretable (for example, linear regression). SHAP is a popular explainability method based on game theory; it produces feature importance scores for each sample. Shapley values represent the average expected marginal contribution of one feature to the prediction after all possible combinations of features have been considered. Permutation importance is another interpretability method producing feature importance scores for each input sample by measuring the change in the model performance (e.g. accuracy) when replacing randomly selected features with noise drawn from the same distribution as the original feature values. The advancements in deep learning brought a vast suite of model-specific interpretability methods [8]. The gradient explanation [9] quantifies the impact on the output of small changes in each input feature and produces a saliency map identifying the input features discriminative of the produced output (here, the DE genes). The “Gradient * Input” [10] method is based on the observation that the saliency maps are enhanced by multiplying the gradient with the input signal.

## 2 Materials and Methods

### Objective

This work compares the performance of traditional DE methods, typically model-agnostic, with that of general ML interpretability models, both model-agnostic and model-specific (to NNs). Existing comparative DE studies [1, 2, 3] are limited to the dedicated DE methods and evaluate their agreement of identified DE genes on annotated cell class assignments (representing the ground truth labels). To incorporate the model-specific analysis, our study introduces a supplementary dimension, representing the clustering model producing the prediction. Three NN-based state-of-the-art clustering methods have been studied: scziDesk [4], scDeepCluster [6] and contrastive-sc [7].

### DE methods

A large experimental study was performed to compare the DE performance of 9 methods. Three statistical tests were employed as a baseline: the t-test, its adaptation to overestimate the variance of each group, and the Wilcoxon rank-sum. These tests were computed between the features of the samples in each of the predicted clusters and the rest of the population. The implementation provided in the scanpy package was used for this exercise. EdgeR and DESingle are included as representative for the methods dedicated to the analysis of scRNA-seq data. All presented approaches return for each predicted cluster of cells an importance score and an adjusted p-value. Next, permutation importance, implemented in the python package ELI5 and SHAP, represent the general approaches to model-agnostic explainability. As there is no implementation for running permutation importance or SHAP on clustering models, a classifier (an XGBoost model) has been trained to learn the predicted clustering and serve as input for the feature importance analysis. Finally, the NN specific explainability models are represented by the gradient and the gradient * input methods. The last four methods return feature importance scores for all samples, characterizing the contribution of each gene to the class assignment. However, as the downstream analysis requires detecting the set of DE genes per cluster, the sample feature importance scores have been averaged per cluster. Our work also studies the impact of applying the DE methods traditionally -directly on the count matrix *D*-or on the preprocessed input data *D*′. *D*′ is obtained by applying the preprocessing phase specific to each of the 3 clustering models to *D*. For brevity, we do not detail here these preprocessing steps, but typical operations are normalization by cell/sample and removal of less variant features. In the remaining manuscript, the experiments corresponding to the traditional setting have been annotated with *.

### Challenges

The absence of “ground truth” information about the DE genes makes the evaluation on real-world datasets is a complex task, usually requiring additional biological experiments for validation. Because the evaluation of real-world datasets is complex and the need for biological experiments would reduce the number of datasets we can analyze, we have limited the scope of this work to the analysis of simulated data, offering complete knowledge about the DE genes in each cluster, as presented in the following section. This simplification allows us to reduce the evaluation strategy to the agreement between the results of different methods and “ground truth” DE gene annotations.

### Evaluation strategy

The DE methods return for each predicted cluster a feature importance score which can be used to sort the genes. The first 5 methods provide an adjusted p-value and employ an arbitrary threshold to rank the genes as DE. Using the same default p-value on various methods produces significant differences in the number of ranked features across methods [3]. The last 4 methods do not provide p-values and require defining a new ranking strategy. A thresholding strategy is proposed to avoid the complexity of aligning ranking strategies. For each cluster, the feature scores are used to sort the genes, apply a threshold to compute the agreement of top 30 and top 100 genes across different methods. As the ground truth annotations also provide the total number of DE genes per cluster, which varies from one datasets/cluster to another but in general is higher than 100, a third evaluation is performed on the complete set of DE genes, denoted as m. Our results report the accuracy at the three ranking levels (e.g. top 30, 100 and m); the lower rankings are employed to determine if the DE genes correctly predicted by a method overlap with the true top DE genes.

### Preprocessing

As cluster labels are permutation tolerant, the cluster labels need to be aligned with the ground truth, prior to the evaluation phase. Existing comparative DE studies analyze datasets with 2 clusters or after classification, which avoids the need for alignment. As a pre-evaluation phase, cluster alignment is performed by choosing the label permutation maximizing the accuracy of the permuted labels.

### Input datasets

As explained in the Challenges paragraph, our analysis was performed only on simulated data. We generated 36 datasets approximating biological scenarios in which we controlled the number of DE (differentially expressed) genes but also the clusters, the samples, the genes and the dropout rates. The DE rate represents how many features (genes) were important, by design, for defining the sample clusters. Knowing the identity of DE genes behind each cluster provides a straight-forward way to validate the results of each DE method executed on top of the clustering predicted by each of the 3 analyzed methods. The dedicated scRNA-seq package *splatter* is used to create balanced and imbalanced datasets (corresponding to uniform and non-uniform distribution of cluster sizes). For each setting, 18 datasets are created with combinations of 2 or 3 clusters, dropout rates of 0.08, 0.17 and 0.3 (dropout.mid parameter) and 0.05, and 0.3 percent of DE genes (de.prob parameter). The cluster size ratios of the im-balanced datasets are 0.5, 0.3 and 0.2. As depicted in Fig. 1ab, dropout impacts more the cluster compactness than the DE rate. Each cluster is annotated with DE positive feature scores; a threshold at log(1.2) is employed to rank the features as DE and create the underlying ground truth; the number of ranked genes per cluster *m* is used as the main evaluation threshold.

**Figure 1:**
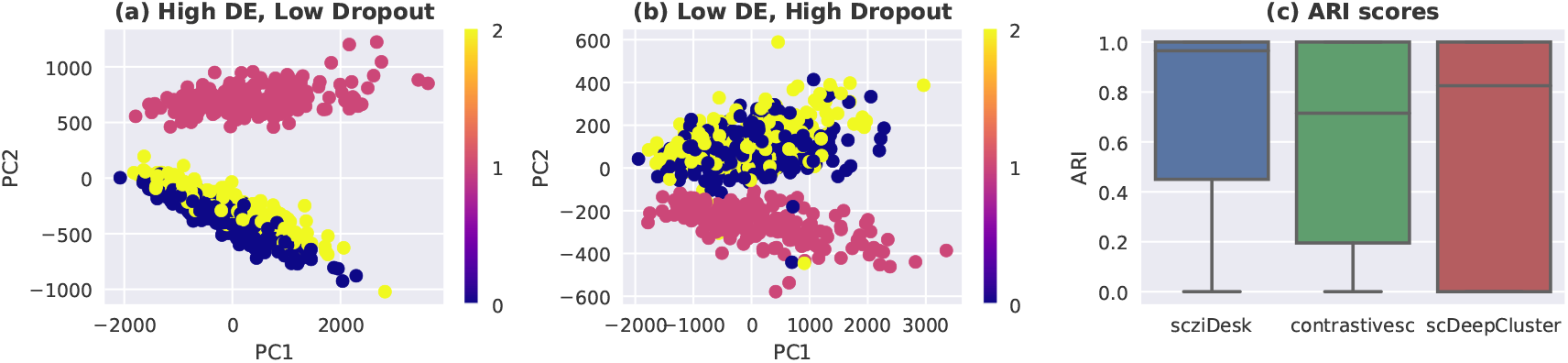
2D PCA representation of datasets having a 0.08 dropout rate and 0.3 DE ratio (panel a) while panel b depicts a dataset with 0.3 dropout rate and 0.1 DE ratio. The colors represent ground truth sample clusters. Panel c depicts the distribution of ARI scores for each of the three clustering methods.

## 3 Results

### Performance

The results depicted in Fig. 2 suggest that the DE methods identify less than 50% of the correct DE genes, both on balanced and imbalanced datasets. The cluster size imbalance doesn’t have a significant impact on the performance of DE models. We leave for future-works the exercise of simulating datasets with more clusters and higher imbalance rates. The gradient method achieved the highest accuracy on the scziDesk and scDeepCluster while on contrastive-sc the results are comparable to the other top performing methods. contrastive-sc employs high levels of NN dropout as data augmentation and thus learns a sparse representation of the input data, penalizing by design the capacity to learn all relevant features and explaining the loss of performance. For disambiguation, NN dropout does not refer to the false zero count events occurring on scRNA-seq data but to a type of layer in artificial neural networks removing (dropping out) an amount of neuron connections to the forward layer. Unlike model-agnostic approaches, the gradients method leverages directly the model which learned to differentiate the clusters, explaining the superior accuracy. We hypothesize that the under-performance of gradient * input is caused by the normalization applied to the input data (here, 0 values do not represent the absence of signal). The worst performing methods are the ML model-agnostic algorithms: feature permutations and SHAP. These techniques were not designed to handle the specificities of scRNA-seq data and, unlike the gradient-based methods, do not have access to the internal state of the model which performed the clustering and learned the discriminating features. The average global accuracy (Fig. 3f) suggests that the best performing method is the gradient saliency and is closely followed by the statistical tests performed on the preprocessed data.

**Figure 2:**
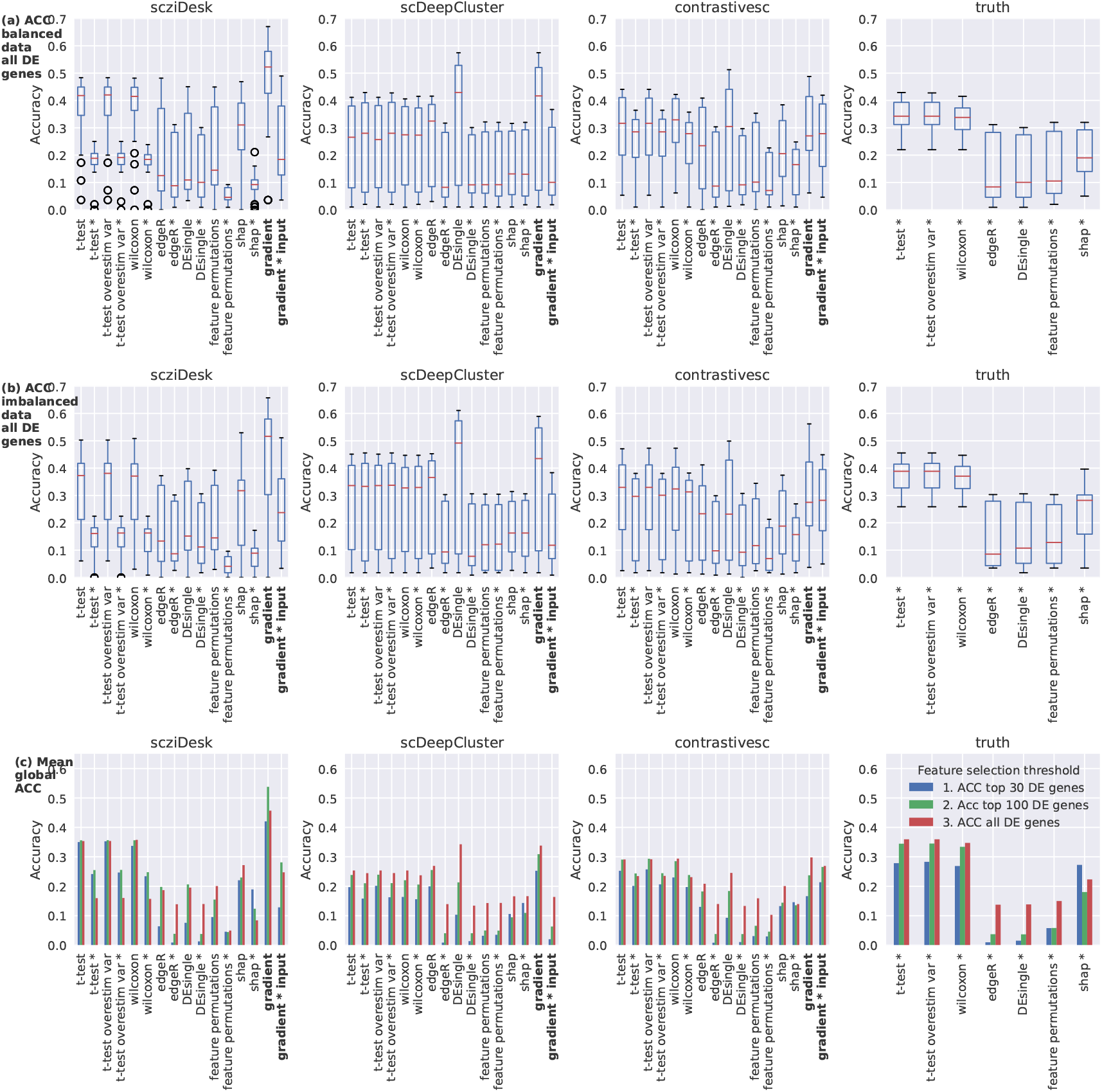
Accuracy of DE methods applied to the clustering predictions made by scziDesk, scDeepCluster and contrastive-sc on the complete set of ground truth features on balanced (panel a) and imbalanced (panel b) datasets. The methods annotated with * were executed on the original count matrix while all other on the data preprocessed by the clustering method (e.g. after gene filtering/normalization). For reference, the last column provides the DE results wrt the ground truth annotations and the entire dataset, replicating other comparative studies. Panel c depicts the average results at different thresholds of accuracy (top 30, top 100 and on all DE genes).

**Figure 3:**
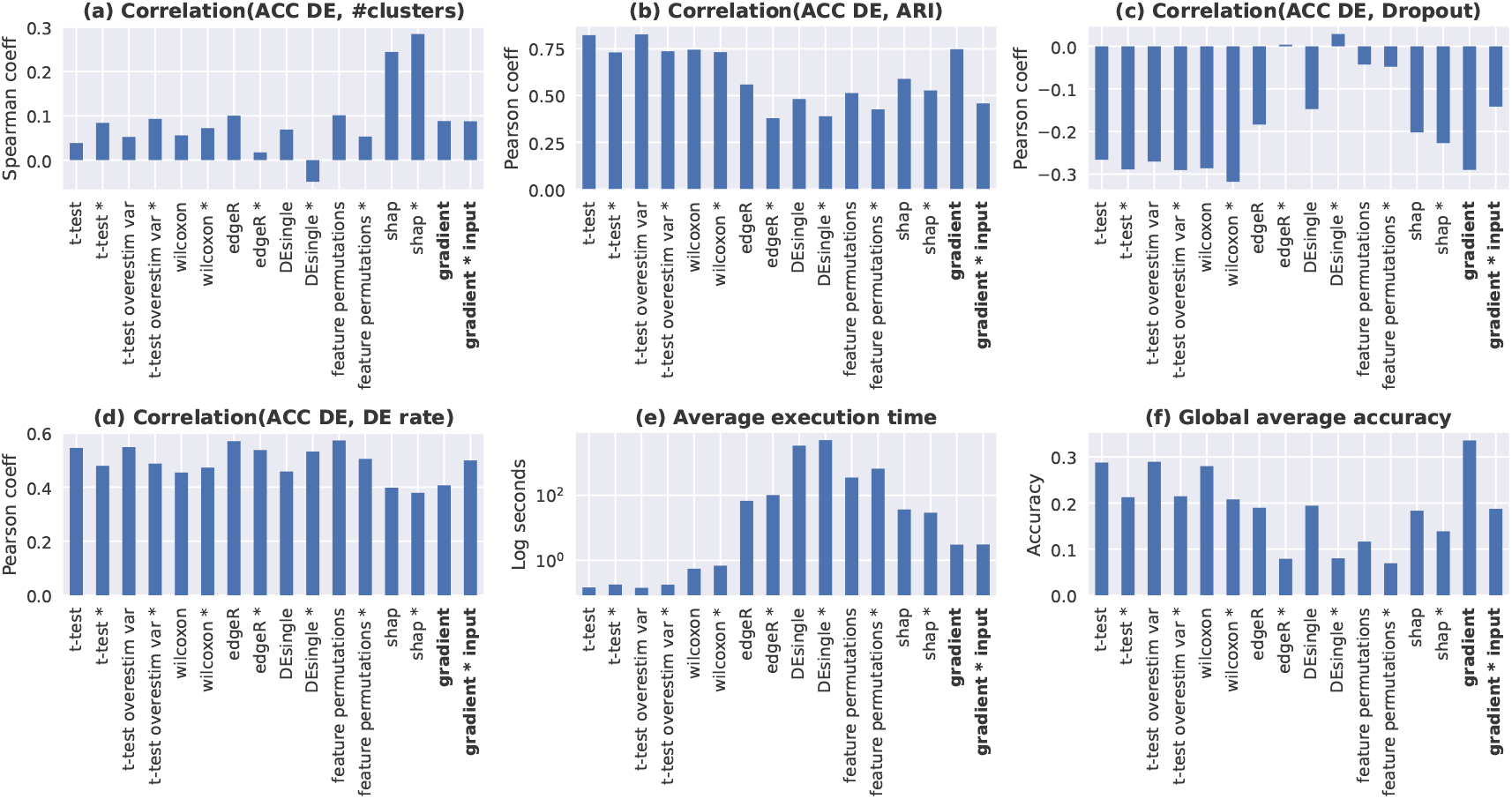
The correlation coefficient between the DE accuracy and the number of clusters (panel a), the clustering ARI score (panel b), the dropout level (panel c) and the DE rate (panel d). Panel e depicts the execution time and panel f the global DE accuracy. Panel a depicts the Spearman correlation while all other panels, the Pearson method. All results are computed across all clustering methods and types of datasets.

### Importance of preprocessing

The preprocessing of clustering methods selects the top variable genes; performing edgeR, DEsingle or statistical tests on this data instead of on the original expression matrix increases the DE accuracy. When using the original counts matrix, the DE methods can still rank as important features which have been discarded and not even used by the model, which confirms the importance of the preprocessing phase. On scDeepCluster, the gradient method and the DEsingle on the preprocessed dataset achieve similar performance. Running the DE analysis on the ground truth produces generally lower results than the gradient methods (but which process partially incorrect predictions Fig. 1c); however, the variability of accuracy scores on the ground truth is significantly lower. Nevertheless, the high Pearson correlation between the clustering ARI scores and the DE accuracy (Fig. 3b) suggest that the improving the clustering results would have a significant impact on the DE accuracy.

### Meta analysis

Our data simulation strategy of exploring the impact of parameters such as various levels of dropout and a number of clusters superior to 2 makes our study complementary to other comparative studies [1, 2, 3]. Increasing the number of clusters in the dataset doesn’t penalize the DE accuracy in general and has a weak correlations (Fig. 3a). Fig. 3c demonstrates that the dropout has a strong negative correlation with the DE accuracy, which can be explained by its negative impact on the clustering performance and thus on the identification of correct clusters. Designed to handle the scRNA-seq dropout, DEsingle is the only method correlating positively the dropout and the DE accuracy. As expected, Fig. 3d demonstrates that the DE accuracy grows with the DE rate in the dataset. All our tests were executed on an Intel(R) Core(TM) i7-9750H CPU @ 2.60GHz with 32 GB RAM. In terms of execution time, the statistical tests and the gradient-based methods are the fastest approaches, requiring on average under 0.5 and 3 seconds respectively to run per dataset. Due to the lower number of genes, the execution time of scziDesk and contrastive-sc is on average under 0.5 while for scDeepCluster under 8 seconds. The most expensive approach is DEsingle (on average one hour per run) followed by feature permutations and SHAP. The repeated sampling performed by these methods explains the underlying computational cost.

### Limitations and future works

The main limitation of this work is the lack of experimentation on real-world datasets, a choice made to provide a simple evaluation strategy. In future works, we foresee extending the scope to real-world analysis as well as to more clustering methods.

## 4 Conclusion

We performed a large comparative study of DE analysis methods. Unlike other existing works, we propose analyzing the results relative to the clustering method and not only to the ground truth. We extended the analyzed methods to model-specific approaches, based on NN gradients. In addition to the traditional DE method, our study included methods typically employed for explaining general machine learning models (feature permutations, SHAP). Using data simulation, we provided complementary results to existing studies, by assessing the impact of dropout and of multiple clusters on the DE analyis. Our results indicate that applying statistical tests on the preprocessed dataset often outperforms the dedicated methods (e.g. DEsingle, edgeR). By showing that gradient-based methods outperform the traditional approaches, while also being significantly faster, we hope that our work will have an impact on developing by design interpretable models, able to both cluster and identify the DE genes. Finally, our research confirms the limited agreement between DE methods also reported in other studies. All code reproducing our experiments is available on [GitHub](https://github.com/ciortanmadalina/sc-interpretability).

